# Abnormal larval neuromuscular junction morphology and physiology in *Drosophila* Prickle isoform mutants with defective axonal transport and adult seizure behavior

**DOI:** 10.1101/2021.12.31.474668

**Authors:** Tristan O’Harrow, Atsushi Ueda, Xiaomin Xing, Salleh Ehaideb, J. Robert Manak, Chun-Fang Wu

**Author notes:** These authors contribute equally (Tristan O’Harrow, Atsushi Ueda). Raymond G. Perelman Center for Cellular and Molecular Therapeutics, The Children’s Hospital of Philadelphia, Philadelphia, PA 19104, USA. Department of Pharmacology, School of Medicine, University of California, Davis, CA, 95616. Experimental Medicine Department, King Abdullah International Medical Research Center, King Saud bin Abdulaziz University for Health Sciences, Ministry of National Guard - Health Affairs, Riyadh, Saudi Arabia (Salleh Ehaideb).

## Abstract

Previous studies have demonstrated that mutations of the *Drosophila* planar cell polarity gene *prickle (pk)* result in altered microtubule-mediated vesicular transport in larval motor axons, as well as adult neuronal circuit hyperexcitability and epileptic behavior. It is also known that mutant alleles of the *prickle*-*prickle* (*pk*^*pk*^) and *prickle*-*spiny-legs* (*pk*^*sple*^) isoforms differ in phenotype but display isoform counterbalancing effects in heteroallelic *pk*^*pk*^*/pk*^*spl*e^ flies to ameliorate adult motor circuit and behavioral hyperexcitability. We have further investigated the larval neuromuscular junction (NMJ) and uncovered robust phenotypes in both *pk*^*pk*^ and *pk*^*sple*^ alleles (heretofore referred to as *pk* and *sple* alleles, respectively), including synaptic terminal overgrowth, as well as irregular motor axon terminal excitability, poor vesicle release synchronicity, and altered efficacy of synaptic transmission. We observed significant increase in whole-cell excitatory junctional potential (EJP) in *pk* homozygotes, which was restored to near WT level in *pk/sple* heterozygotes. We further examined motor terminal excitability sustained by presynaptic Ca^2+^ channels, under the condition of pharmacological blockade of Na^+^ and K^+^ channel function. Such manipulation revealed extreme Ca^2+^ channel-dependent nerve terminal excitability in both *pk* and *sple* mutants. However, when combined in *pk/sple* heterozygotes, such terminal hyper-excitability was restored to nearly normal. Focal recording from individual synaptic boutons revealed asynchronous vesicle release in both *pk* and *sple* homozygotes, which nevertheless persisted in *pk/sple* heterozygotes without indications of isoform counter-balancing effects. Similarly, the overgrowth at NMJs was not compensated in *pk/sple* heterozygotes, exhibiting an extremity comparable to that in *pk* and *sple* homozygotes. Our observations uncovered differential roles of the *pk* and *sple* isoforms and their distinct interactions in the various structural and functional aspects of the larval NMJ and adult neural circuits.

## Introduction

Mutations of the *Drosophila* planar cell polarity gene *prickle* (*pk*) alter cuticular bristles and ommatidia organization (Gubb & García-Bellido, 1982; Tree et al., 2002). Two Pk protein isoforms Pk^pk^ and Pk^sple^ are expressed post-embryonically and disruption of either isoform leads to different aspects of bristle defects (Gubb et al., 1999). Additionally, in larval motor axons, defective microtubule dynamics and vesicular transport have been reported in an isoform-specific fashion (Ehaideb et al., 2014). In *pk*^*pk*^*/+* mutant larvae, axonal vesicular transport is decreased while *pk*^*sple*^/+ mutants show enhanced anterograde vesicle transport (heretofore referred to as *pk/+* and *sple/+* alleles, respectively). Moreover, overexpression of the Pk^sple^ isoform in WT motor neurons (which mimics the Pk^pk^ reduction relative to Pk^sple^ in *pk/+* mutants) results in reversal of ∼30% of the microtubules in the axons, while overexpression of the Pk^pk^ isoform (which mimics the Pk^sple^ reduction relative to Pk^pk^ in *sple*/+ mutants) results in altered microtubule dynamics, and thus increased microtubule plus-end polymerization.

Notably, the *pk* gene is conserved across humans, mice, zebrafish, and *Drosophila*, and has been linked to epileptic behaviors in all these organisms (Tao et al., 2011). Adult *sple1*/+ flies show ataxia, hypersensitivity to mechanical shock (bang sensitivity), and a myoclonic-like seizure phenotype, mirroring the epilepsy syndrome observed in human patients with *PRICKLE* gene mutations (Ehaideb et al., 2014, 2016). Interestingly, bang-sensitive and seizure phenotypes are greatly suppressed in *sple1*/*pk1* heterozygotes. Furthermore, manipulating gene dosage by transgenic overexpression of Pk isoforms can mimic the seizure phenotype of *sple1*/+, indicating a counterbalancing interaction between the Pk and Sple isoforms. In addition, the *sple1*/+ epileptic phenotypes can be rescued by a mutation of the motor protein subunit Kinesin light chain (*Klc)*, suggesting an association of epileptic phenotypes to neuronal microtubule-mediated transport (Ehaideb et al., 2016).

To help fill in the gap between the molecular defects and the behavioral phenotypes during development and function of motor synapses, we examined cellular physiological and morphological defects in *pk1* and *sple1* mutant alleles, using the same larval neuromuscular preparation (Jan & Jan, 1976; Wu et al., 1978; Atwood et al., 1993), in which the axonal transport defects were observed. Pre-synaptic axonal terminal morphology and post-synaptic subsynaptic reticulum (SSR) structures can be readily visualized with immunohistochemical staining against Horse Radish Peroxidase (HRP) and Disks-large (Dlg), respectively (Jan & Jan, 1982; Budnik et al., 1996). Synaptic transmission at the NMJ can be analyzed by intracellular recording of excitatory junctional potentials (EJPs) for examination of whole-cell response (Jan & Jan, 1976) or by focal extracellular recordings to resolve individual bouton responses (Kurdyak et al., 1994; Renger et al., 2000; Xing & Wu, 2018a, 2018b). This approach allows direct association of axonal transport defects (Ehaideb et al., 2014) with observed morphological and physiological phenotypes in *pk1* and *sple1* mutant larvae.

We found defective nerve-terminal morphology and synaptic transmission in *pk1* and *sple1* alleles. Alterations in NMJ growth, synaptic efficacy, nerve terminal excitability and synaptic vesicle release synchronicity were demonstrated, but their severity was not simply in parallel to the adult phenotypes associated with the various heteroallelic combinations among *pk*^+^, *pk1, sple*^*+*^, and *sple1* (Ehaideb et al., 2014; 2016). Further studies are required to establish in detail the functional roles of Pk and Sple isoforms linked to specific aspects of neuronal form and function in different cell types and neural circuits, at the various developmental stages.

## Materials and Methods

### Fly stocks

Flies and larvae were housed in bottles containing a cornmeal-agar-based medium (Frankel & Brousseau, 1968). Flies were reared at a room temperature of 22 ± 1°C. The *pk*^*pk1*^, *pk*^*pk30*^, and *pk*^*sple1*^ mutations were introgressed into the control wild-type Oregon-R (OR) line for a minimum of 6 generations to reduce the possibility of genetic background effects, and all these lines used in this study have been documented previously (Ehaideb et al., 2014). The whole-cell EJP data for *pk*^*pk30*^ were consistent with *pk*^*pk1*^ and therefore they were pooled. Heterozygous mutants (*pk*^*pk1*^/+ & *pk*^*sple1*^/+) were generated by crossing the *pk* mutant to OR. Heteroallelic mutants (*pk*^*pk1*^/*pk*^*sple1*^) were generated by cross between the two mutant lines.

### Immunohistochemistry and NMJ morphology

Wandering third instar larvae were dissected in HL 3.1 saline (Feng et al., 2004) containing 0 mM Ca^2+^. Larval preparations were fixed in 3.7% formaldehyde in Ca^2+^-free HL 3.1 for 30-40 minutes. Preparations were stained with goat anti-HRP conjugated to FITC (Jackson ImmunoResearch Laboratories, West Grove, PA, USA) (1:50 dilution) at 4°C for 12-48 h. For double-staining with anti-HRP and anti-Dlg, monoclonal AB 4F3 (Parnas et al., 2001) against Dlg was used (Developmental Studies Hybridoma Bank (DSHB), University of Iowa, Iowa City, IA, USA). Fixed larval preps were incubated with the primary monoclonal AB 4F3 at 1:100 in phosphate buffer saline, at 4°C for 12-48 h, rinsed, and further incubated in phosphate buffer saline containing anti-HRP (conj. FITC) and a secondary TRITC-conjugated anti-mouse IgG antibody (Jackson ImmunoResearch Laboratories, West Grove, PA, USA), both at 1:50. Larval NMJ images were gathered on a Leica DM IL LED microscope and a Leica TCS SP8 confocal microscope (Leica Microsystems Inc., Buffalo Grove, IL, USA), using the Leica Application Suite X software.

Examination of larval NMJ morphology was carried out on type Ib boutons in muscle 4 (M4) in abdominal segments A2, A4, and A6. Data from different segments was consistent within each genotype, and were pooled for statistics. For counting of NMJ branches, a branch was defined as a terminal process carrying at least two synaptic boutons (Zhong et al., 1992).

### Larval neuromuscular electrophysiology

All electrophysiological recordings were performed in HL3.1 saline (Feng et al., 2004) with the composition (mM): 70 NaCl, 5 KCl, 4 MgCl_2_, 10 NaHCO_3_, 5 trehalose, 115 sucrose, 5 HEPES (pH adjusted to 7.1). For nerve-evoked whole-cell EJP recording, 0.2 mM CaCl_2_ was added to the Ca^2+^-free HL 3.1 saline, and for extracellular focal recording and whole-cell “plateau-like” potential recoding, 0.1 mM CaCl_2_ was added.

Intracellular recording of excitatory junction potentials (EJPs) was adapted from Ueda and Wu (2006). To evoke EJPs triggered by nerve action potentials, the segmental nerve was severed from the ventral ganglion, and 0.1-ms stimuli were applied to the cut end of the nerve through a suction pipette. Stimulation amplitude was adjusted to 2 times threshold voltage to ensure uniform stimulation conditions among experiments. To enable analysis of presynaptic terminal excitability, we also performed electrotonic stimulation on the NMJ (Wu et al., 1978; Ganetzky & Wu, 1982; 1983). Briefly, tetrodotoxin (TTX, 3µM) was applied to block Na^+^ channels. To achieve direct electrotonic stimulation of the terminal, a longer duration (2-ms) stimulus was applied near the hemisegment entry point by drawing in the segmental nerve to the suction pipette, so as to effectively apply different levels of depolarization with passive electrotonic spreading to the terminal (Lee et al., 2014). In this manner, synaptic terminal CaV channels were directly triggered by local depolarization, independent from invasion of axonal Na^+^ action potentials (Wu et al., 1978). For the generation of plateau-like potentials, multiple K^+^ channel species, including *Shaker* (Kv1), *Shab* (Kv2) and *eag* (Kv10) (Jan et al., 1977; Ganetzky & Wu, 1982; 1983; Ueda & Wu, 2009), were blocked by co-application of 4-aminopyridine (4-AP) and tetraethylammonium (TEA) to allow the development of full-blown regenerative Ca^2+^-action potentials that sustain prolonged transmitter release (Ueda & Wu, 2006; 2009; Lee et al., 2014).

The method for loose-patch extracellular focal recording has been described previously (Kurdyak et al., 1994; Renger et al., 2000; Xing & Wu, 2018a), based on the early works on the frog and crayfish MNJs (Fatt and Katz, 1952; Dudel and Kuffler, 1961). Briefly, A glass electrode was chipped at the tip, polished (Microforge de Fonbrune, Series A, No. 310, Aloe Scientific, St. Louis, MO, USA) to an opening 10-15 μm wide (covering one to three boutons), and the shank of the electrode was bent (for perpendicular access to the muscle surface with the polished opening) with heat from the coil of the Microforge. The electrode contained an AgCl-Ag wire, and was filled with HL3.1 (0.1 mM Ca^2+^). The electrode opening was positioned on top of a distal portion of the type Ib NMJ of larval muscle 4 of abdominals segments A3-A6. Local synaptic transmission signals were collected through an AC amplifier with a low- and high-frequency cutoff of 0.1 Hz and 50 kHz, and passed through the NEUROCCD-SM256 system into a PC computer running the NeuroPlex application (RedShirt Imaging LLC, Decatur, GA, USA).

Chemical compounds were purchased from the following companies: TTX from Alomone Labs (JBP, Israel), 4-AP and HEPES free acid from Sigma (St. Louis, MO USA), TEA and trehalose from Across Organics (Geel, Belgium), NaCl and KCl from RPI (Mount Prospect, IL USA), CaCl_2_-2H_2_O, MgCl_2_-6H_2_O, and NaHCO_3_ were purchased from Thermo Fisher Scientific (Waltham, MA USA).

### Statistics

One-way ANOVA and F-test with sequential Bonferroni correction were performed as described in the corresponding section of Results.

## Results

We found that mutant *pk* larvae displayed abnormal NMJ morphology compared to WT controls (Figure 1A, B). As summarized in Figure 1C, homozygotes for the *pk1* and *sple1* mutations showed increases in both the numbers of axonal terminal branches and synaptic boutons, with axonal terminal branches defined as a process carrying 2 or more boutons (cf. Zhong et al., 1992). Larvae with one copy of each isoform mutation (p*k1/sple1*) also displayed greater numbers of boutons than WT. Further, one copy of either mutation (*pk1/*+ or *sple1/*+) was able to confer an increase in synaptic boutons comparable to that observed in the homozygous mutants (Figure 1B and Table 1). However, a suggestive trend of increase in branch number was not statistically significant compared to wild-type (Table 1). This indicates semi-dominance of the synaptic bouton phenotype of the larval NMJ. A characteristic morphological phenotype particular to the *pk1* larval NMJ is the appearance of long branches without typical structures of boutons, not encountered in other genotypes examined here (Figure 1B). As revealed by anti-HRP and anti-Dlg staining, these branches lack both presynaptic bouton-like enlargements and the postsynaptic subsynaptic reticulum. (Figure 1A, B). In contrast, anti-HRP and anti-Dlg staining indicate that *sple1* larvae were prone to exhibiting an opposite phenotype, with boutons packed closely together along the terminal branch (Figure 1A, B).

**Figure 1.**
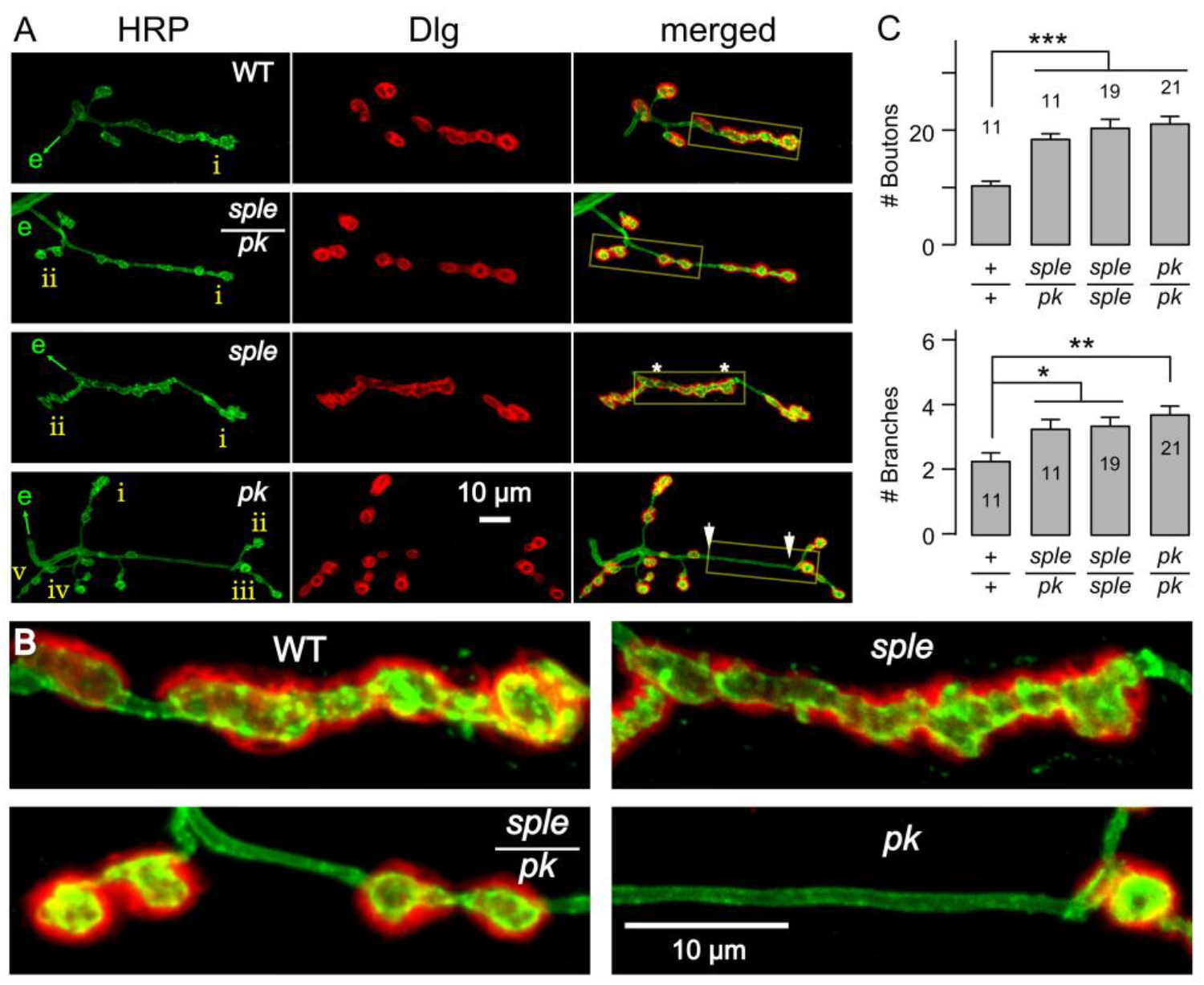
(A) NMJ morphology of Prickle mutants *pk*^*pk1*^/*pk*^*pk1*^and *pk*^*sple1*^/*pk*^*sple1*^, the *pk*^*pk1*^/*pk*^*sple1*^ double heterozygote, and WT. Presynaptic membrane is stained with anti-HRP, and subsynaptic reticulum is stained with anti-Dlg (monoclonal AB 4F3) (see methods). Nerve entry is indicated by e with a green arrow. Yellow boxed region is expanded in B. Note complex boutons structures are indicated by anti HRP antibody staining in sple. Between some of the *pk*’s boutons were long interbouton areas (arrow heads). (B) Expanded confocal images of boutons in A. (C) Increased numbers of boutons and branches in Prickle mutants. Error bars = SEM. Number of NMJs are indicated. *, **, and *** indicates significant differences (*p* < 0.05, 0.01, and 0.001, respectively with one-way ANOVA).

**Table 1.** Comparison among heterozygous *pk*^*sple1*^/+, *pk*^*pk1*^/+, and *pk*^*sple1*^/*pk*^*pk1*^. Mean ± SEM (n) is shown. * indicates statistically significant difference from *pk*^*sple1*^/*pk*^*pk1*^ (*p*< 0.05, one-way ANOVA). Note that most parameters were not significantly different among these three genotypes.

| Genotype | # boutons<br>per NMJ | # branches<br>per NMJ | Fraction of stimuli with asynchronous release |  |
| --- | --- | --- | --- | --- |
|  |  |  | 1 Hz | 10 Hz |
| *sple/+* | 17.9 ± 0.7 (21) | 2.2 ± 0.2 (21) * | 0.14 ± 0.05 (8) | 0.27 ± 0.09 (5) |
| *pk/+* | 19.7 ± 0.8 (21) | 2.7 ± 0.3 (21) | 0.36 ± 0.09 (10) | 0.50 ± 0.14 (5) |
| *sple/pk* | 18.4 ± 1.0 (11) | 3.3 ± 0.3 (11) | 0.33 ± 0.06 (10) | 0.31 ± 0.05 (10) |

We further examined the efficacy of nerve action potential-triggered synaptic transmission with intracellular recording of postsynaptic EJP, the whole-cell muscle response to transmitter release from the entire synaptic terminal arbors. EJP amplitude was increased in the *pk1* mutant compared to WT control (Figure 2A, B), which could be simply a consequence of a larger amount of presynaptic vesicle release from the increased number of synaptic boutons in *pk1*. However, no such EJP amplitude increase was observed in the *sple1* mutant, even though it also displayed greater numbers of boutons than WT. This suggests a more complex relationship between number of synaptic boutons and EJP size in mutants of the two Pk isoforms. A potential mechanism may involve differential efficacy in synaptic vesicle release from *pk* and *sple* boutons, as well as differences in their postsynaptic response. Interestingly, muscle input resistance was skewed toward higher values in both *pk* and *sple* compared to WT (WT = 5.9 ± 1.4 MΩ, n=8; *pk* = 8.0 ± 1.1 MΩ, n=10; *sple* = 7.7 ± 3.4 MΩ, n=11; mean ± SD; *p* < 0.01, t-test for *pk*; *p* < 0.05, F-test for *sple*). Since EJP size increase occurred only in *pk* but not in *sple*, muscle input resistance changes could not be the sole reason for changes in EJP size.

**Figure 2.**
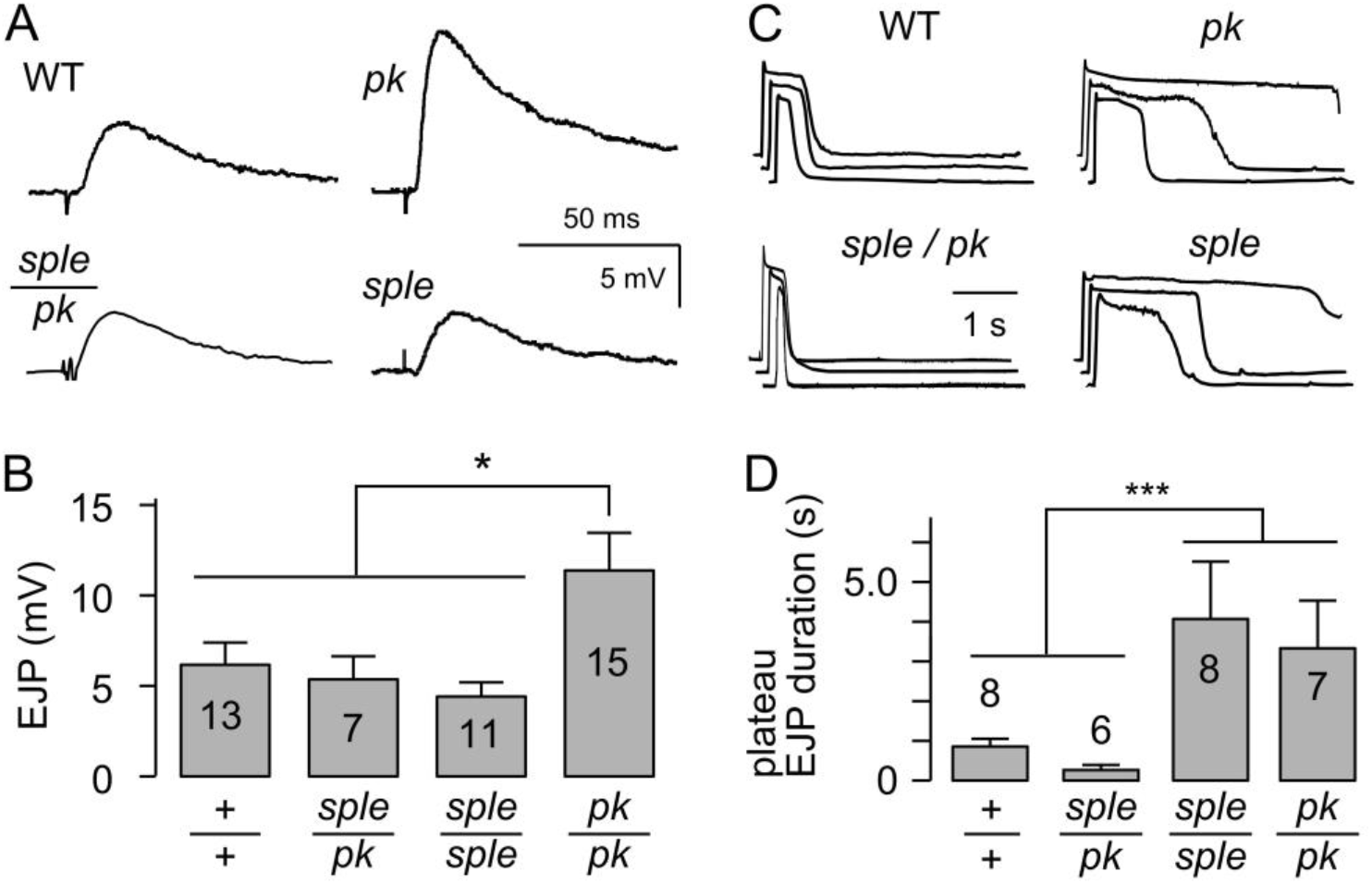
Abnormal synaptic transmission in *pk*. (A, B) Increased EJP size in *pk*^*pk1*^/*pk*^*pk1*^. Whole-cell EJPs were measured in HL3.1 saline containing 0.2 mM Ca^2+^. # of NMJs are indicated inside the bars. Error bars = SEM. * indicates significant differences (*p* < 0.05, One-way ANOVA). (C) Altered Ca^2+^-dependent excitability in *pk*^*pk1*^/*pk*^*pk1*^ mutant motor axon terminals. Motor axon terminals were directly activated by electrotonic stimulation in the presence of K^+^ channel blockers (4-AP, 200 μM and TEA, 20 mM) and a Na^+^ channel blocker TTX. Under this condition, prolonged “plateau” EJPs supported by continuous transmitter release were recorded (see Results & Discussion). Amplitude is normalized to steady-state. (D) The durations of plateau EJPs were prolonged and more variable in *pk*^*sple1*^/*pk*^*sple1*^ and *pk*^*pk1*^/*pk*^*pk1*^. *pk*^*pk1*^/*pk*^*sple1*^ resembled WT. *** indicates significant differences among the group (*p* < 0.001, F-test with sequential Bonferroni correction). Error bars = SEM. # of NMJs are indicated.

Prompted by the previously reported counterbalancing effect of *pk* and *sple* mutations on hyperexcitability and epileptic phenotypes in adult *pk1*/*sple1* heterozygotes. (Ehaideb et al., 2014), we investigated how this genetic interaction is implicated in the context of larval NMJ physiology and morphology. First, we found that EJP amplitude in p*k1/sple1* was indeed similar to WT and *sple1*, an apparent counter balancing action of *sple1* of suppressing *pk1*-induced excessive transmission (Figure 2A, B). However, muscle input resistance was increased also in *pk*/*sple* (10.2 ± 4.5 MΩ, n=7; *p* < 0.01, F-test), suggesting that whole-cell synaptic current in *pk*/*sple* might be decreased compared to WT.

The driving force for synaptic current appeared normal in all the above mutants since muscle resting membrane potential was within a range similar to WT (−64 ∼ -69 mV, on average). Therefore, the EJP remediation in *pk1*/*sple1* larvae might be a consequence of differential effects of *pk1* and *sple1* on synaptic release mechanisms or the growth mechanisms of synaptic boutons. Nevertheless, morphologically no counter balancing effects were observed since the numbers of synaptic boutons and branches remained comparable to either *pk1*/*pk1* or *sple1*/*sple1*, significantly higher than WT control (Figure 1). Therefore, one should examine other components of synaptic transmission mechanisms for the *pk*-*sple* interactions in determining transmission efficacy, since the driving force and the synaptic bouton numbers could not account for the observed EJP size changes.

Excitability control of synaptic terminals is crucial for the synaptic vesicle release process and involve interplay among various Na^+^, Ca^2+^, and K^+^ channels. Under normal physiological conditions, synaptic transmission is initiated by axonal action potential invasion and presynaptic Ca^2+^ influx to trigger vesicular release for subsequent opening of postsynaptic glutamate receptor channels. By using the Na_V_ channel blocker tetrodotoxin (TTX) to remove the participation of Na_V_ in the control of synaptic transmission, transmitter release can still be induced by direct electrotonic stimulation of the presynaptic terminal to evoke depolarization of controlled duration and amplitude (Ganetzky &Wu, 1982; Ueda& Wu, 2009). Thus, we were able to investigate potential alterations in nerve terminal excitability that sustains Ca^2+^ infux for neurotransmitter release in isolation of axonal Na^+^ AP invasion.

Some subtle defects in presynaptic nerve terminal excitability could be further revealed using this approach with further blockade of repolarizing currents through K^+^ channels. We used the A-type K^+^ channel blocker 4-AP and a broad-spectrum K^+^ channel blocker TEA (Peng & Wu, 2007; Singh & Wu, 1999) to investigate the extent Ca^2+^ influx through the presynaptic Ca^2+^ channels could maintain presynaptic terminal depolarization, in order to sustain prolonged transmitter release. Under this extreme condition, sustained synaptic transmission, from hundreds of ms to seconds, manifested as prolonged postsynaptic depolarization or “plateau EJP” could be readily recorded intracellularly in the postsynaptic muscle fiber (Ganetzky & Wu, 1982; Ueda & Wu, 2006). By using this approach, we found that both *pk1* and *sple1* displayed plateau EJPs of longer duration than WT, while the double heterozygote of *pk1/sple1* resembled WT (Figure 2C,D).

These results suggest that disruptions of either Pk or Sple isoform alters Ca^2+^-driven excitability, indicating potential alterations in Ca^2+^-channel properties or reduced sensitivity of certain K^+^ channels to pharmacological blockade. Nevertheless, the changes in Pk and Sple isoforms, when combined in heterozygotes, create a compensatory effect to restore Ca^2+^-driven synaptic terminal excitability.

To uncover potential alterations in the synaptic vesicle release process in individual boutons, we used the extracellular focal recording technique (see methods) to capture postsynaptic current dynamics associated with presynaptic release. Focal recording allows us to examine quantal events of single vesicular release at fast time resolution, enabling detection of altered release kinetics, such as the synchronicity or temporal precision in the release process (Renger et al., 2000; Ueda & Wu, 2009). We found that in the presence of TTX, focal synaptic transmission evoked by electrotonic stimulus was frequently followed by one or more asynchronous releases in the *pk1* and *sple1* mutant larvae, indicating altered regulation of the vesicular release mechanism (Figure 3A). These asynchronous releases appeared most strikingly in homozygotes of *pk1* and *sple1* mutants (*pk1*/*pk1* and *sple1*/*sple1*), and to a lesser extent in heterozygotes (*pk1*/*+* and *sple1*/+) (Figure 3A,B, Table 1). Asynchronous transmissions were more clearly revealed by higher frequency repetitive stimuli (10 Hz), although these events were also seen even at 1 Hz. Notably, this general pattern is true for *pk1/sple1* heterozygous (Figure 3B, Table 1). There is no indication of a compensatory interaction between the mutant Pk and Sple isoforms in the temporal precision regulation of the release process.

**Figure 3.**
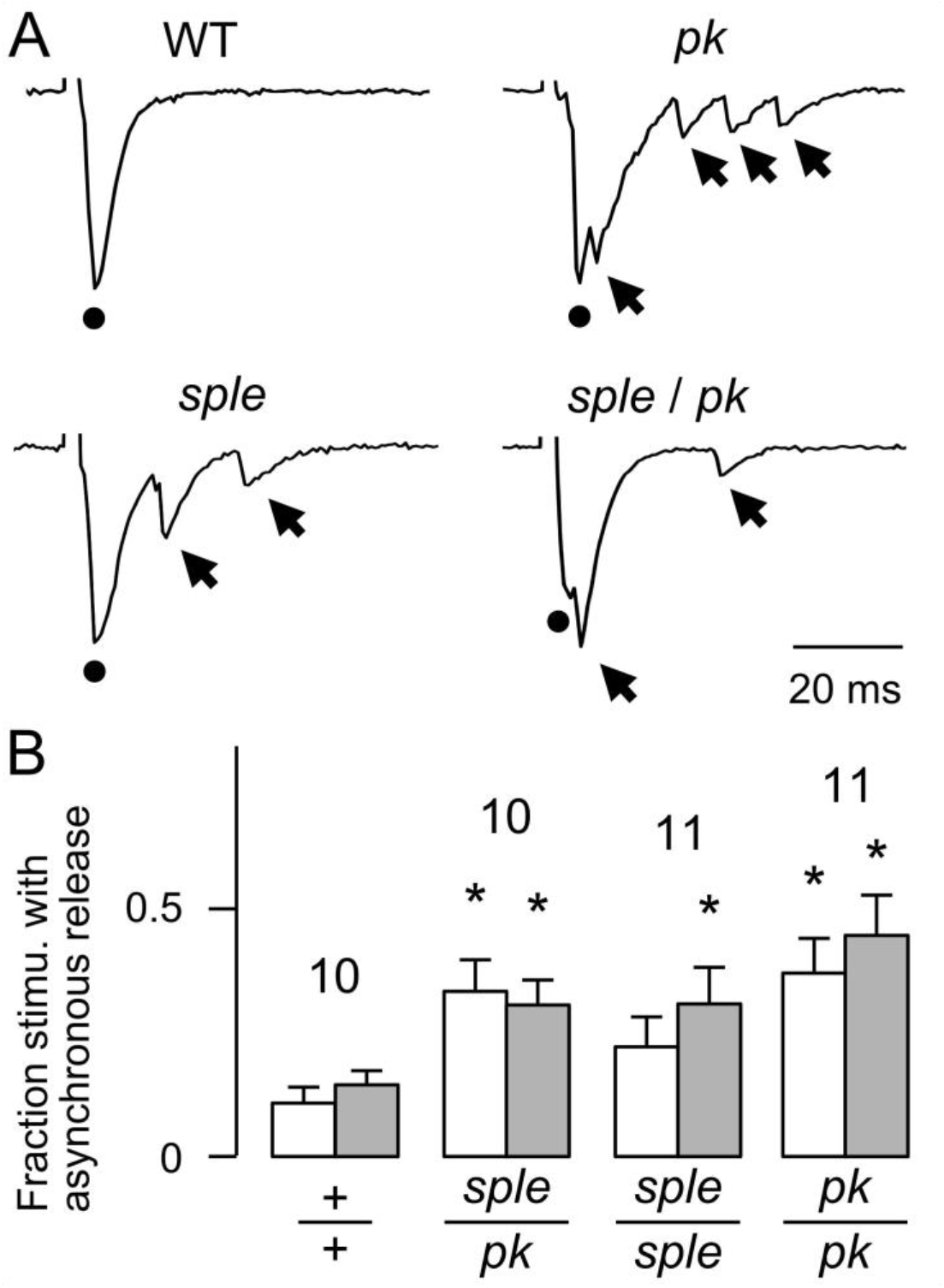
(A) Focal EJPs (fEJP) evoked by electrotonic stimulation of nerve terminals (2 ms 10Hz) in the presence of an Na_V_ blocker (TTX, 3 μM). *pk*^*sple1*^/*pk*^*sple1*^, *pk*^*pk1*^/*pk*^*pk1*^, and the *pk*^*pk1*^/*pk*^*sple1*^ double heterozygote exhibited asynchronous neurotransmitter releases after stimulation. Dots indicate the evoked synchronous release. Arrows indicate asynchronous releases. Recoded from type Ib boutons on muscle 4. (B) Proportion of evoked fEJPs followed by one or more asynchronous releases within 100 ms after each stimulus. Stimuli were applied at 1 Hz (10 s train, white bars) and 10 Hz (2 s train, gray bars). Stimulus voltage was adjusted to approx. two-times the threshold for release. * indicates significant deviation from WT (*p* < 0.05; one-way ANOVA). Error bars = SEM. # of NMJs are indicated.

## Discussion

The *prickle* gene in *Drosophila* has been best characterized for its role in promoting the proper polarity of bristles in cuticle structures (Gubb & García-Bellido, 1982; Gubb et al. 1999), which involves intracellular vesicular or organelle transport. Similarly, Homologous Pk mutations in vertebrates are known to affect surface structures of epithelial origin, including hairs and cilia (Sowers et al., 2014), in addition to neurological phenotypes, such as epilepsy (Tao et al., 2011; Todd & Bassuk, 2018; Ehaideb et al. 2014; 2016). Therefore, the *Drosophila* larval NMJ can be further studied to explore a wider range of cellular phenotypes in order to gain a deeper understanding and to uncover novel common mechanisms underlying defects in the various cell types of different tissues.

Detailed mutational effects of *pk1* and *sple1* have been described in prior studies for adult flies (Ehaideb et al., 2014; 2016), including motor circuit hyperexcitability and epilepsy-like behavioral phenotypes. However, such defects have been linked to *sple1* but nearly absent in *pk1 flies*. Further, these phenotypes were suppressed in *pk1/sple1* heterozygotes, indicating a counterbalancing effect of mutations of the two isoforms (Ehaideb et al., 2014). In the same report, different aspects of defective vesicular axonal transport were demonstrated in both *pk1/+* and *sple1/+* larval motor neurons (Ehaideb et al., 2014).

We have now examined, in the same larval NMJ preparation, synaptic structure and function to investigate potential consequences of altered axonal transport. We confirmed that both *pk1* and *sple1* exhibited dominant phenotypes of synaptic terminal growth and synaptic release control (Table 1). Significantly, we observed increased size of nerve-evoked EJPs in *pk1* but not in *sple1* (Figure 2A). However, in both *pk1* and *sple1* mutant larvae, we discovered striking phenotypes of NMJ overgrowth (Figure 1), asynchronous vesicular release (Figure 3), and enhanced Ca^2+^-dependent terminal excitability leading to plateaued EJPs (Figure 2B). We found *pk1*/*sple1* counterbalancing rescue only for abnormal EJPs (nerve-evoked and electrotonically induced, Figure 2A & B), while there was no indication of the counterbalancing effects for nerve terminal overgrowth (Figure 1) and asynchronous vesicle release (Figure 3) in the heterozygotes.

It should be noted that the results from the prior studies do not necessarily contradict with the current findings. The seizure studies carried out in adult flies employed an electroconvulsive stimulation (ECS) paradigm, where high-frequency intense stimulation to the brain triggered seizure-like neural circuit activity, manifested as motor output of action potential bursts recorded in flight muscles (Ehaideb et al., 2014; 2016). In contrast, the study presented here focused on the cellular phenotypes at the larval NMJ, encompassing motor axon properties and synaptic transmission examined in different subcellular compartments. Such results may have some bearing of overall cellular phenotypes in mutants of *pk* and *sple* isoforms but do not provide sufficient information directly relevant to the control mechanisms underlying the various excitability patterns at the circuit and system levels. Conceivably, synaptic mechanisms revealed in the larval motor axon and presynaptic terminal in *pk1* and *sple1* mutants can take part, but only in some restricted aspects, in the excitability control of the central circuits that contribute to the seizure phenotype. A more complete picture could involve various types of interneurons and transmitter systems beyond the glutamatergic motor neuron described here.

It will be desirable to establish in future studies whether adult NMJs innervated by glutamatergic motor neurons display corresponding defects, in order to support or refute the possibility that increased neuronal excitability in *pk1* is restricted to the larval stage, and circuit-level hyperexcitability and seizure predisposition in *sple1* only arise upon metamorphosis to the adult stage. Similarly, the synaptic structure of the adult *pk* or *sple* mutant has not been documented and it will be important to investigate whether the current findings of altered larval NMJ morphology (Figure 1) can be extended to the homologous NMJs in the adult stage. If so, an immediate question would be whether the mutant overgrowth phenotype could be generalized to some non-glutamatergic synapses, e.g. cholinergic, GABAergic, dopaminergic, octopaminergic, serotonergic, etc.

Previous studies have also shown that adult circuit hyperexcitability and epileptic behavior linked to *sple1* can be suppressed by appropriately balancing the isoform (*pk1* and *sple1*) dosage (Ehaideb et al., 2014). Counterbalancing effects were also found in *pk1/sple1* larval NMJ in mitigation of increased EJP size due to *pk1* (Figure 2A,B). Hetrozygous *pk1/sple1* larvae also lacked the increased Ca^2+^ excitability of the nerve terminal in individual *pk1* and *sple1* mutants (Figure 2C,D). These results indicate that the counterbalancing effect of Prickle isoforms observed in adult flies is also present in some aspects of the larval motor neuron function. However, the features of larval NMJ overgrowth and asynchronous vesicular release appeared to be independent from isoform dosage. Thus, the roles of Prickle isoforms in regulating motor neuron morphology and physiology may involve interacting partners different from those governing motor circuit excitability and seizure predisposition.

Pk mutations confer a spectrum of neurological phenotypes most likely through interactions with a variety of synaptic proteins. It has been implicated that Synapsin interacts with Pk, leading to alterations in neuronal morphology and physiology. Immunochemical evidence indicates that mouse PRICKLE1 protein can be physically associated with the protein SYNAPSIN I, a major synaptic phosphoprotein, and that in the *Drosophila* larval NMJ transgenic Prickle also co-localizes with Synapsin (Paemka et al., 2013). It is known that *Drosophila* Synapsin influences synaptic development, synaptic vesicle formation, and transmitter release (Vasin et al., 2014). It will be important to examine whether physical association between Synapsin and Prickle proteins plays a role in generation of the NMJ morphology and physiology characteristic of *prickle* mutants observed in this work. Notably, in addition to its association with epilepsy, a link has been implicated between human *PRICKLE* mutations to autism (Sowers et al., 2013; Todd & Bassuk, 2019), which may involve disrupted physical association between PRICKLE1 and SYNAPSIN I (Paemka et al., 2013).

It is evident that the *Drosophila* larval neuromuscular preparation will remain as a highly relevant model to investigate the complex role of Prickle in neural development and function. Further studies on genetic interactions between *pk* with genes encoding additional synaptic proteins in the various cellular processes, e. g. intracellular vesicular trafficking in regulating transmitter vesicle release and recycling, and membrane expansion and retraction during synaptic growth, to highlight a potentially critical role of Pk in synaptic development, function and disease.

## Acknowledgement

We thank Atulya Iyengar for discussion, Lydia Luton for fly stock maintenance. This work has been supported by USPHS NIH grants AG051513 (AU, XX, TO, C-FW) and NS098590 (CFW, RJM). SE was supported by a fellowship from King Abdullah International Medical Research Center, Riyadh, Saudi Arabia.

